# Amputation Triggers Long-Range Epidermal Permeability Changes in Evolutionarily Distant Regenerative Organisms

**DOI:** 10.1101/2024.08.29.610385

**Authors:** Kelly E. Dooling, Ryan T. Kim, Elane M. Kim, Erica Chen, Adnan Abouelela, Benjamin J. Tajer, Noah J. Lopez, Julia C. Paoli, Connor J. Powell, Anna G. Luong, S.Y. Celeste Wu, Kara N. Thornton, Hani D. Singer, Aaron M. Savage, Joel Bateman, Tia DiTommaso, Duygu Payzin-Dogru, Jessica L. Whited

## Abstract

Previous studies have reported that amputation invokes body-wide responses in regenerative organisms, but most have not examined the implications of these changes beyond the region of tissue regrowth. Specifically, long-range epidermal responses to amputation are largely uncharacterized, with research on amputation-induced epidermal responses in regenerative organisms traditionally being restricted to the wound site. Here, we investigate the effect of amputation on long-range epidermal permeability in two evolutionarily distant, regenerative organisms: axolotls and planarians. We find that amputation triggers a long-range increase in epidermal permeability in axolotls, accompanied by a long-range epidermal downregulation in MAPK signaling. Additionally, we provide functional evidence that pharmacologically inhibiting MAPK signaling in regenerating planarians increases long-range epidermal permeability. These findings advance our knowledge of body-wide changes due to amputation in regenerative organisms and warrant further study on whether epidermal permeability dysregulation in the context of amputation may lead to pathology in both regenerative and non-regenerative organisms.

**GRAPHICAL ABSTRACT:** 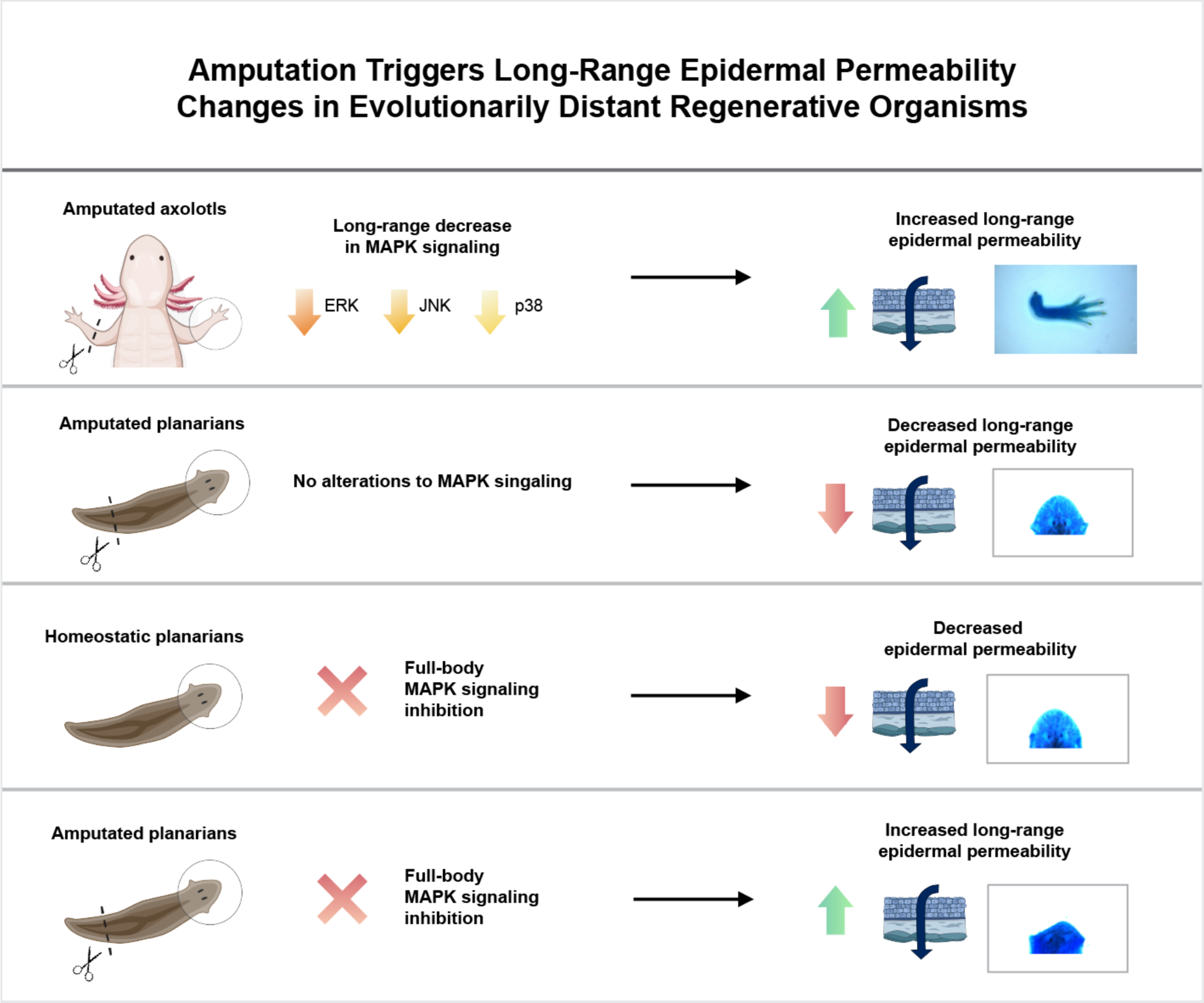

## INTRODUCTION

Limb amputation is a common treatment for life-threatening complications of vascular disease, diabetes, and trauma, with 150,000 Americans per year undergoing a major amputation (Dillingham et al. 2005; Ziegler-Graham et al. 2008). Amputees often experience poor postoperative life expectancies (reviewed in Qaarie 2023), which may stem from dysregulated body-wide responses to amputation. For example, amputation-induced stress signaling is thought to accelerate pathological features of diabetes and cardiovascular disease (Modan et al. 1998; Peles et al. 1995). Wound healing in amputees is a complex process with frequent dermatologic complications such as infection, which can become systemic (Berger et al. 2023; Heim et al. 2003; reviewed in Pascale and Potter 2014). As such, investigating the whole-body consequences of amputation is clinically important for addressing morbidity among amputees.

Indeed, in animal models, mounting evidence indicates that amputation triggers body-wide responses that extend beyond the wound site (reviewed in Ricci and Srivastava 2018; Sun and Poss 2023). Planarian and acoel worm amputation (Baguñà 1976; Fan et al. 2023; Saló and Baguñà 1984; Srivastava et al. 2014; Wenemoser and Reddien 2010), sea star arm amputation (Hernroth et al. 2010), and mouse injury (Rodgers et al. 2017; Rodgers et al. 2014) all trigger long-range responses to injury. In axolotls (*Ambystoma mexicanum*), amputation induces a body-wide cell cycle reentry (Johnson et al. 2018; Payzin-Dogru et al. 2024 preprint), termed “systemic activation,” which primes the animal for faster future regeneration: the remaining limbs of previously amputated, systemically activated animals regenerate faster than the limbs of animals naïve to amputation (Payzin-Dogru et al. 2024 preprint), and the systemic activation response leads to body-wide transcriptional changes in tissues, including the epidermis (Payzin-Dogru et al. 2024 preprint).

In salamanders, the epidermis has traditionally been studied at the amputation plane in the context of epimorphic regeneration. A first step of amphibian regeneration is the formation of a specialized wound epidermis—a proliferative layer of keratinocyte progenitors—over the injury site (reviewed in Han et al. 2005; Satoh et al. 2008). The wound closure phase is rapid, featuring the migration of epidermal cells across the amputation plane (Ferris et al. 2010), with full wound closure by 6 to 8 hours after the initial injury (Lévesque et al. 2010; Satoh et al. 2008). In contrast, mammalian re-epithelialization and wound healing occur over multiple days (reviewed in Han et al. 2005), a process characterized by scar tissue deposition (reviewed in Takeo et al. 2015). The salamander wound epidermis is transcriptionally distinct from homeostatic epidermis (Knapp et al. 2013), and the subsequent coalescence of progenitor cells into a visible blastema—a zone of proliferating cells that differentiate into a regenerated limb— requires this specialized wound epidermis (Goss 1956; reviewed in McCusker et al. 2015; Mescher 1976). Salamander skin regeneration after excisional injury also features rapid wound closure and avoids fibrosis (Seifert et al. 2012).

Planarians are evolutionarily distant to vertebrates, but also exhibit robust regeneration responses (reviewed in Sánchez Alvarado 2012), during which the wound epidermis is thought to play a role in regulating blastema formation (Scimone et al. 2022). Re-epithelialization at the injury site features rapid migration of and structural changes in epidermal cells, with coverage of the injury site by 24 hours post injury (Morita and Best 1974; Pedersen 1976). Planarian regeneration is driven by the proliferation of pluripotent cells called neoblasts (reviewed in Ge et al. 2022; Newmark and Sánchez Alvarado 2000; Wagner et al. 2011), which give rise to new epidermal cells (Morita and Best 1974; Pedersen 1976; Wurtzel et al. 2017) at defined differentiation stages (Tu et al. 2015; van Wolfswinkel et al. 2014; Wurtzel et al. 2017).

While the regulatory roles of the wound epidermis have been widely studied during regeneration, long-range epidermal responses to amputation beyond the wound site, specifically those that lead to functional changes, are largely uncharacterized. Here, we investigate epidermal tissues distant to the amputation plane in axolotls and planarians and find distinct changes in epidermal permeability after amputation in these organisms. We report that amputation triggers a long-range increase in epidermal permeability in axolotls, accompanied by a long-range epidermal downregulation of MAPK signaling. Additionally, pharmacologically inhibiting MAPK signaling in regenerating planarians increases long-range epidermal permeability. These findings suggest evolutionarily-conserved, body-wide epidermal responses to amputation, highlighting the importance of examining long-range epidermal permeability and barrier dysregulation, which may lead to pathology, in the context of amputation and multi-tissue injury.

## RESULTS

### Epidermis in limbs contralateral to amputation is more permeable than that of homeostatic and regenerating limbs

Previous studies report body-wide changes in the axolotl epidermis after amputation, including increased cell cycle reentry in epidermal cells (Johnson et al. 2018; Payzin-Dogru et al. 2024 preprint) and robust transcriptional changes in skin-resident epidermal and small secretory cell populations (Payzin-Dogru et al. 2024 preprint). Here, we investigate whether there is a functional difference between the epidermis of homeostatic (naïve) and regenerating axolotls, examining how amputation alters the ability of the epidermis to act as a barrier.

To test this, we performed a toluidine blue stain—a technique used to assess epidermal permeability in mice (Cameron et al. 2007; Hardman et al. 1998; Indra and Leid 2011; Koch et al. 2000; Nakajima et al. 2007; reviewed in Schmitz et al. 2015) and leaf cuticles (Tanaka et al. 2003; Zhao et al. 2019)—on homeostatic limbs, limbs contralateral to the originally amputated limb (contralateral limbs), and regenerating limbs in axolotls (Fig. 1A-B). We found that contralateral limbs were significantly darker than homeostatic limbs at 14 days post amputation (dpa) (p adj = 0.0001), 21 dpa (p adj = 0.0073), and 70 dpa (p adj < 0.00001), indicating an increase in epidermal permeability in contralateral limbs at these time points (Fig. 1C). We also found significant differences between contralateral limbs at different time points during regeneration, with contralateral limbs displaying a general increase in permeability over the 70- day regeneration period (Fig. S1A).

**Fig 1.**
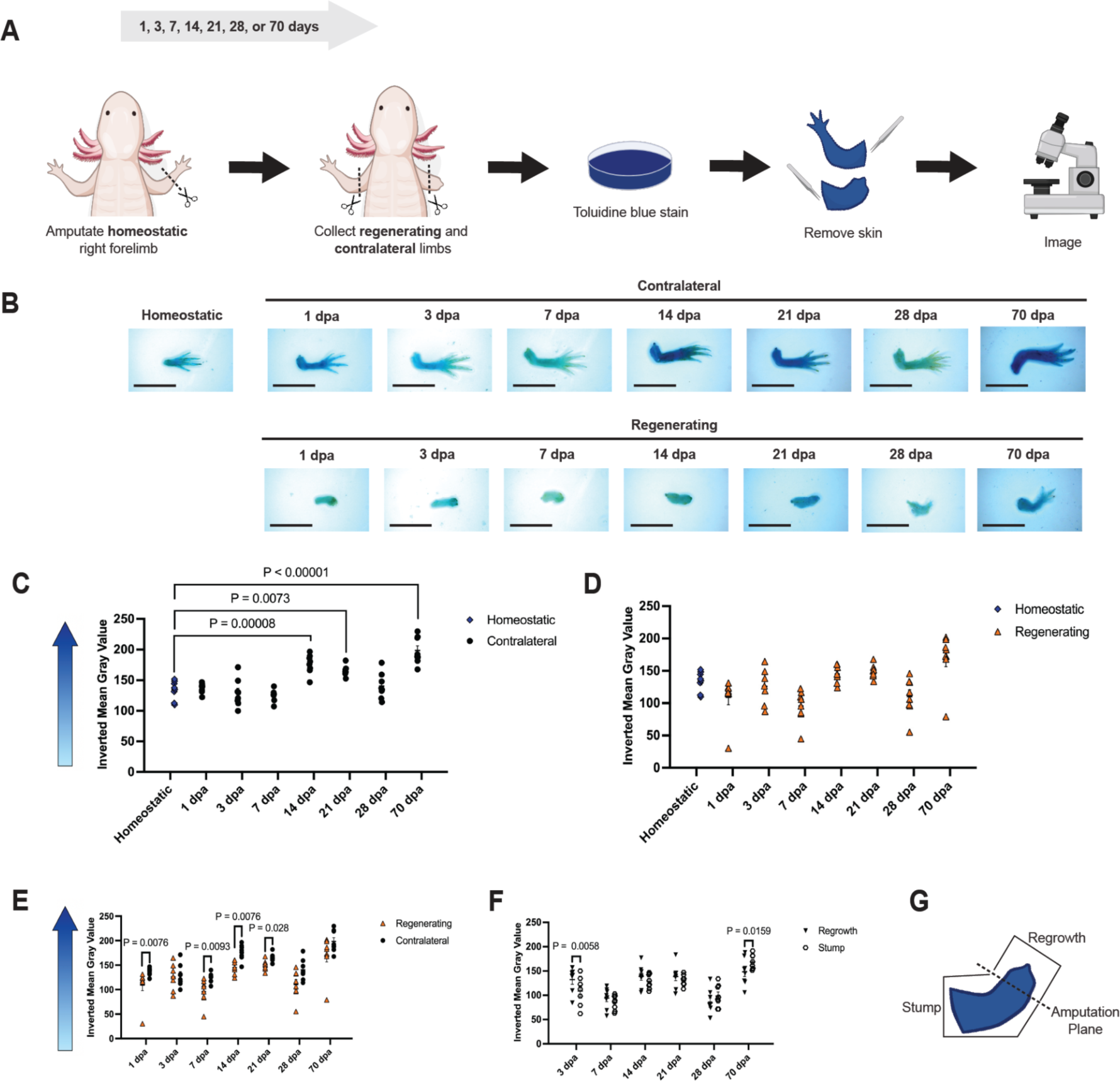
Epidermis in limbs contralateral to amputation is more permeable than that of homeostatic and regenerating limbs. (A) Experimental set-up. (B) Representative images of stained limbs. Scale bars are 1 cm. (C-F) Quantification of epidermal permeability, as measured by inverted mean gray values, with higher inverted mean gray values indicating darker limbs and increased epidermal permeability. Quantification of (C) homeostatic vs. contralateral, (D) homeostatic vs. regenerating, and (E) regenerating vs. contralateral limb stains. (F) Quantification of the regrown vs. original (stump) areas of regenerating limbs, with (G) region specification. All data points represent biological replicates. Data is shown as mean ± standard error. Statistical significance was determined using one-way ANOVA with Tukey’s HSD (C), Kruskal-Wallis ANOVA with Dunn’s test of multiple comparisons (D), Mann-Whitney U tests with Holm correction (E), and paired two-way ANOVA with Sidak correction (F).

Our assay indicated no significant differences in epidermal permeability between homeostatic and regenerating limbs at any time point during regeneration. However, a general increase in epidermal permeability was observed during regeneration (Fig. 1D, S1B). We observed the most rapid increase in permeability during the late stages of blastema formation and proliferation (7-14 dpa, Fig. S1B). These results suggest an overall increase in epidermal permeability in regenerating limbs starting at 3 dpa and continuing throughout the entire 70-day regeneration period (Fig. S1B).

When we compared contralateral to regenerating limbs, we found that contralateral limbs were significantly darker than regenerating limbs at multiple time points—1 dpa (p adj = 0.0076), 7 dpa (p adj = 0.0093), 14 dpa (p adj = 0.0076), and 21 dpa (p adj = 0.0280)—observing that contralateral limbs are initially more permeable than regenerating limbs, but this difference is absent at full regeneration (Fig. 1E). Regrowth (defined in Fig. 1G) in regenerating limbs was significantly darker than stump tissue proximal to the amputation plane at 3 dpa (p adj = 0.0058, Fig. 1F). However, stump tissue (defined in Fig. 1G) was significantly darker than regrowth in regenerating limbs at 70 dpa (p adj = 0.0159, Fig. 1F).

### Homeostatic and contralateral epidermis do not differ in nuclear density or thickness

Migrating epidermal cells at the wound surface of adult eastern newts (*Notophthalmus viridescens*) are known to differ in shape and exhibit increased intercellular separation compared to homeostatic epidermal cells (Repesh and Oberpriller 1980); however, long-range morphological changes in the epidermis after amputation are unknown. We assessed whether there were gross morphological changes in the epidermis of contralateral limbs in axolotls, which might be driving amputation-induced, long-range increases in epidermal permeability.

We performed hematoxylin and eosin (H&E) staining to visualize epidermal morphology in homeostatic epidermis compared to contralateral epidermis at 1, 3, 7, 14, and 21 dpa (Fig. 2A), and found no significant differences in either epidermal thickness or nuclear density between groups, averaged across three epidermal regions (defined as radial, carpal, and digit epidermis, Fig. 2B-D). Region-specific (Fig. 2B) comparisons also yielded no significant differences in epidermal thickness or nuclear density between homeostatic and contralateral epidermis at the given time points (Fig. 2E-F).

**Fig 2.**
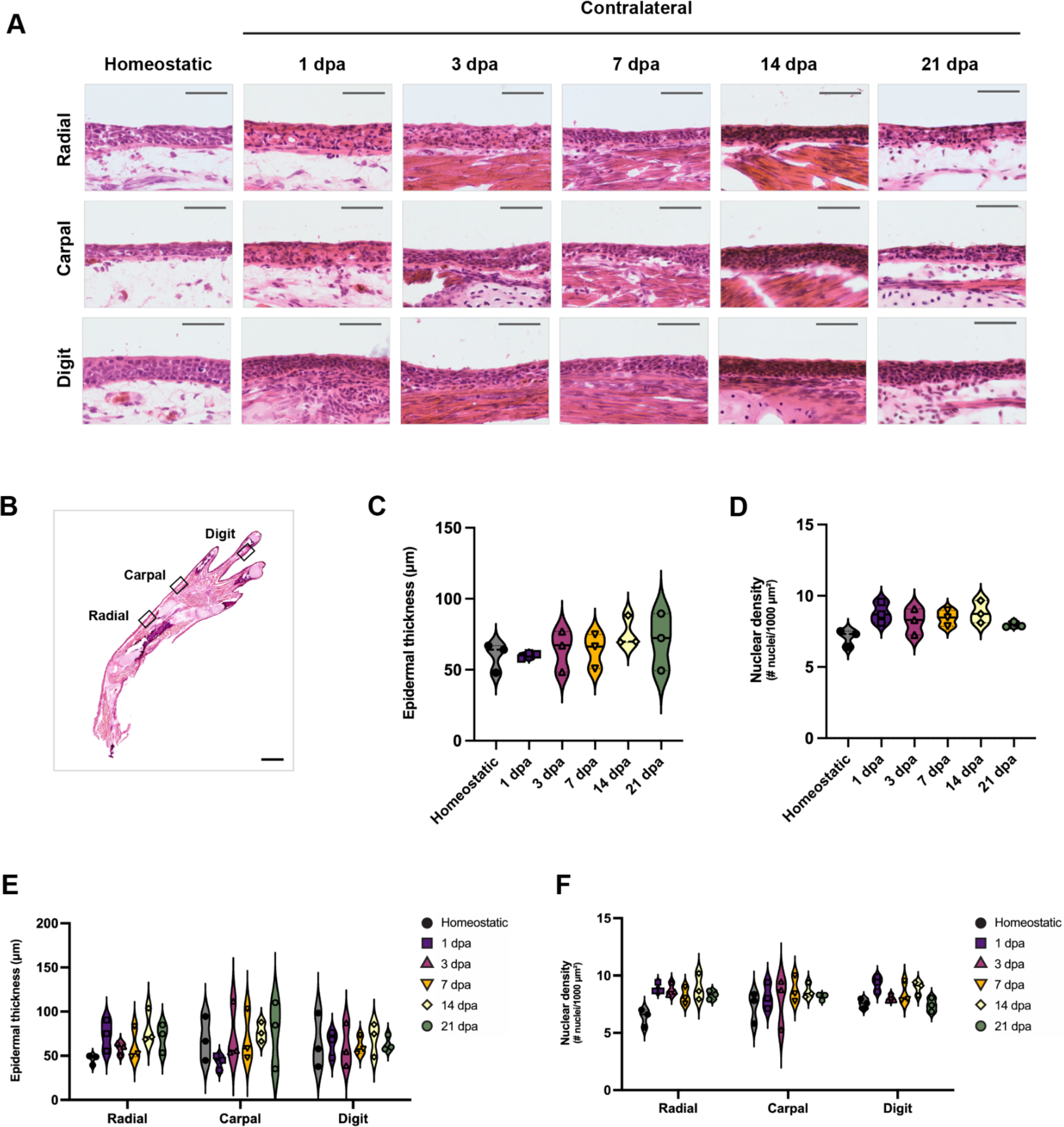
Homeostatic and contralateral epidermis do not differ in nuclear density or thickness. (A) Representative images of hematoxylin and eosin stains of homeostatic and contralateral limbs at 1 to 21 dpa. (B) Segmenting methodology for quantification, whereby 800 μm segments of radial, carpal, and digit epidermis were selected for quantification. (C-D) Quantification of (C) epidermal thickness and (D) nuclear density, averaged across all three regions. (E-F) Quantification of (E) epidermal thickness and (F) nuclear density, separated by region. Scale bars are (A) 100 μm or (B) 1 mm. All data points represent biological replicates. Statistical significance was determined using ordinary one-way ANOVA with Tukey correction (C, D), paired two-way ANOVA with Sidak correction (E), or Kruskal-Wallis ANOVA with Dunn’s correction (F).

### Claudin 1 and collagen IV protein expression is consistent between homeostatic and contralateral epidermis

Since the dysregulation of epidermal tight junctions and the basement membrane can both cause epidermal barrier dysfunction in other contexts, such as mammalian species (reviewed in Breitkreutz et al. 2013; Furuse et al. 2002), we assessed the expression of two structural proteins in homeostatic and contralateral axolotl epidermis: (1) claudin 1, a principal component of epidermal tight junctions, and (2) collagen IV, a protein expressed in the epidermal basement membrane.

When comparing homeostatic and contralateral epidermis at 3 and 7 dpa, we found no significant differences in claudin 1 expression (Fig. 3A-D). We also found no significant differences in collagen IV expression between homeostatic and 10 dpa contralateral epidermis (Fig. 3E-H). Region-specific (radial, carpal, and digit regions, Fig. 3B, F) comparisons between homeostatic and contralateral epidermis similarly yielded no significant differences in either claudin 1 or collagen IV expression (Fig. 3D, H).

**Fig 3.**
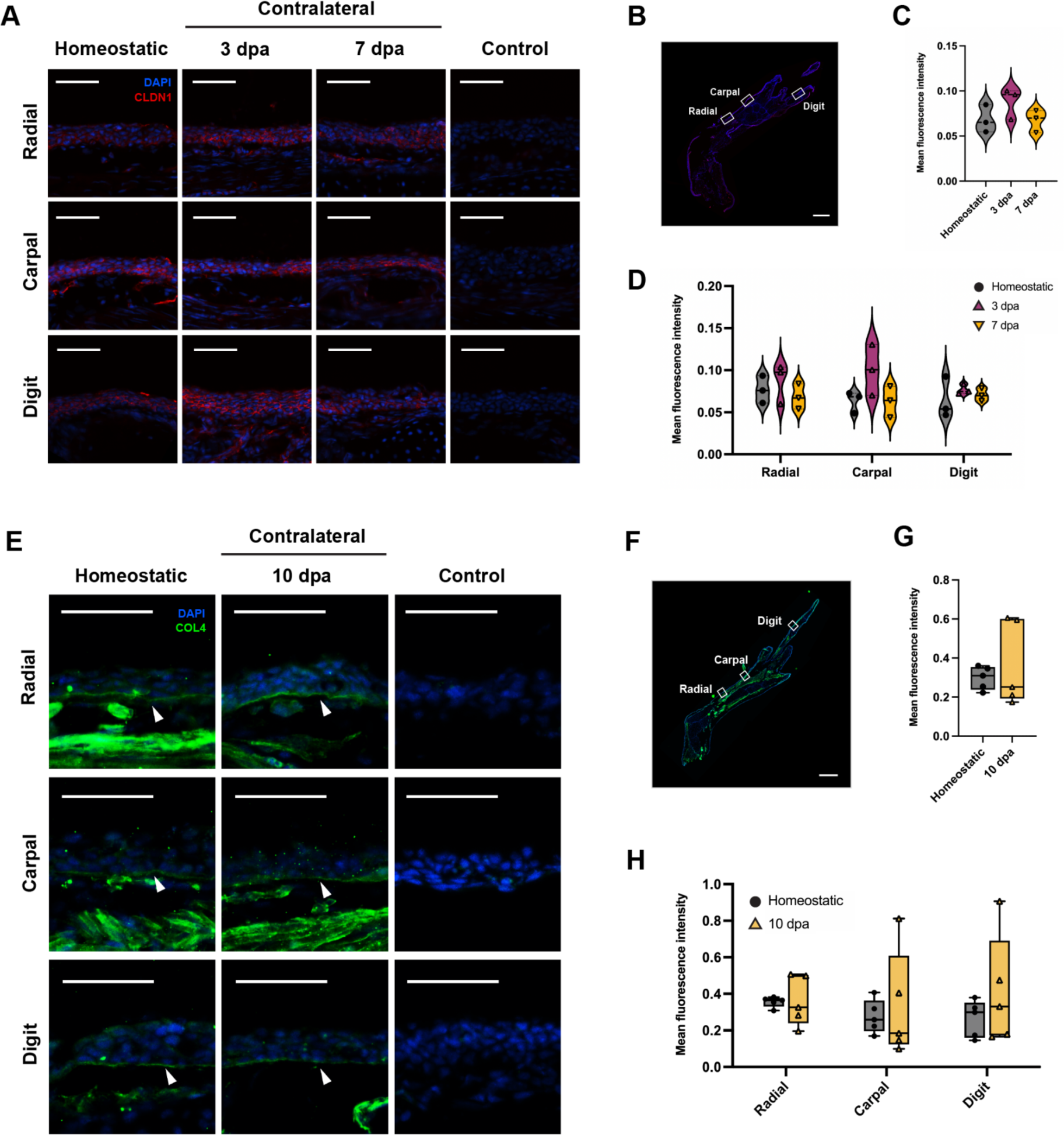
Claudin 1 and collagen IV protein expression is consistent between homeostatic and contralateral epidermis. (A) Representative images of epidermal claudin 1 immunofluorescence including negative control. (B) 800 µm segmenting methodology for radial, carpal, and digit epidermal quantification. (C-D) Claudin 1 quantification, (C) averaged across all three regions, and (D) separated by region. (E) Representative images of epidermal collagen IV immunofluorescence, including negative control. Arrowheads highlight the basement membrane. (F) 300 µm segmenting methodology for radial, carpal, and digit epidermal quantification. (G-H) Collagen IV quantification, (G) averaged across all three regions, and (H) separated by region. Scales bars are (A, E) 100 μm or (B, F) 1 mm. All data points represent biological replicates. Statistical significance was determined using ordinary one-way ANOVA with Tukey correction (C), paired two-way ANOVA with Sidak correction (D, H), or two-tailed paired t-test (G).

### MAPK signaling is downregulated in contralateral epidermis

Dysregulated mitogen-activated protein kinase (MAPK) signaling impairs epidermal barrier repair in mouse and human skin models (Kanemaru et al. 2017; Koehler et al. 2011), prompting us to investigate whether regulation of MAPK signaling is linked to increased epidermal permeability in axolotls. MAPK signaling is upregulated in the growth zone at early stages of axolotl tissue regeneration (Franklin et al. 2017; Ohashi et al. 2021; Rao et al. 2009; Sabin et al. 2015; Sader et al. 2019; reviewed in Sader and Roy 2022; Yun et al. 2014); however, MAPK signaling responses in tissues distant from the amputation plane have yet to be characterized.

We found that the epidermis of contralateral limbs exhibits a robust downregulation in MAPK signaling compared to the epidermis of homeostatic limbs naïve to amputation. Western blot analysis indicated significant decreases in extracellular signal-regulated kinase (ERK) phosphorylation in contralateral epidermis at 1 dpa (p adj = 0.0028), 3 dpa (p adj = 0.0012), and 7 dpa (p adj = 0.0015), each compared to homeostatic epidermis (Fig. 4A-B). We also observed significant decreases in Jun N-terminal kinase (JNK) phosphorylation in contralateral epidermis at 1, 3, and 7 dpa (all p adj < 0.0001, Fig. 4E-F). Although differences in p38 phosphorylation between homeostatic and contralateral epidermis were insignificant, contralateral epidermis at 3 dpa (p adj = 0.0718) and 7 dpa (p adj = 0.0845) exhibited a strong trend of decreased p38 phosphorylation, compared to homeostatic epidermis (Fig. 4C-D).

**Fig. 4.**
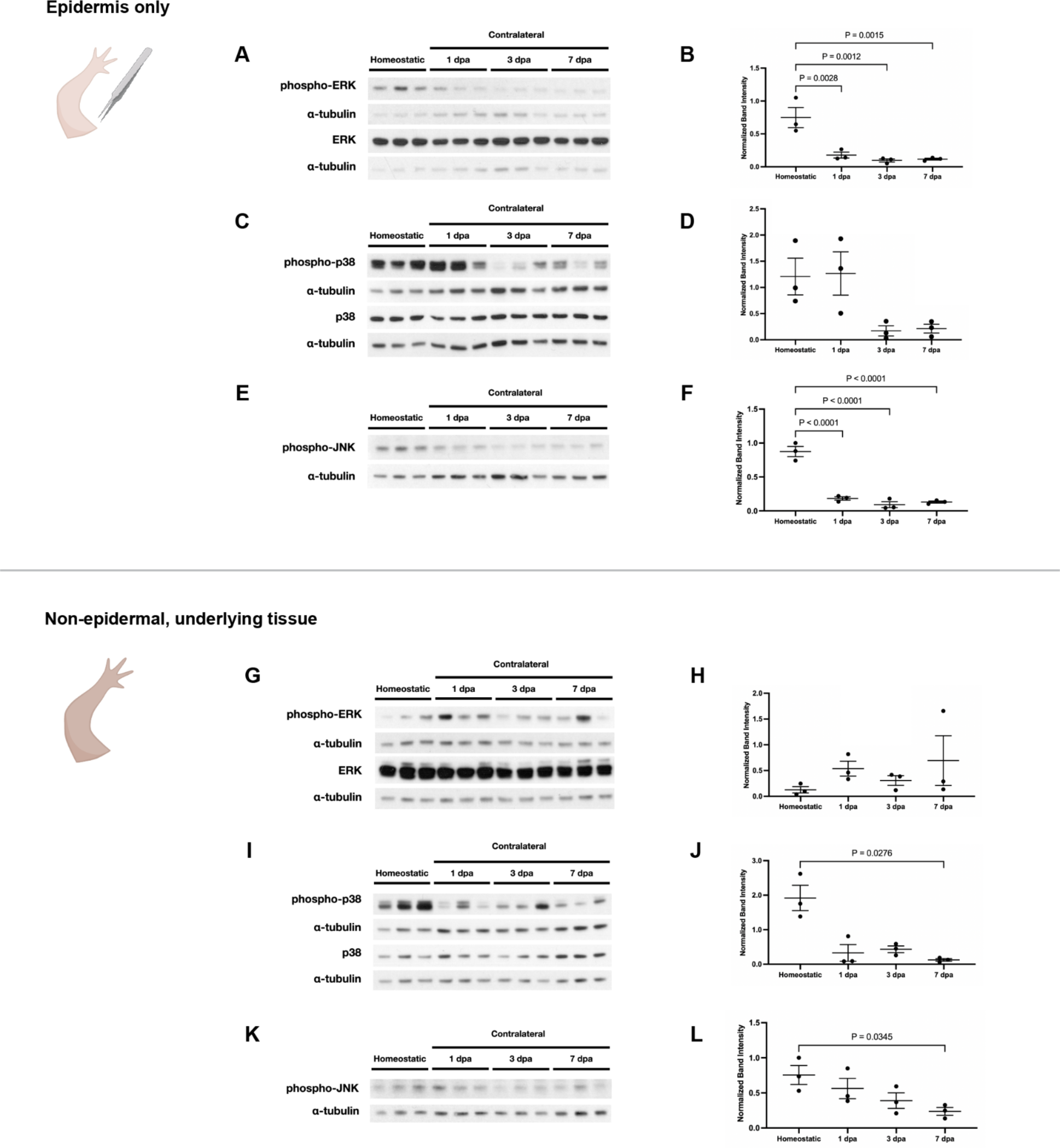
MAPK signaling is downregulated in contralateral epidermis. (A, C, E, G, I, K) Western blots assessing ERK, p38, and JNK phosphorylation in (A, C, E) epidermal and (G, I, K) underlying non-epidermal tissue from homeostatic and contralateral limbs, with (B, D, F, H, J, L) quantifications. Band intensity was generated from the ratio of phosphorylated to total protein, measured by area under curve (AUC) values, normalized within groups and to alpha-tubulin AUC values. (F, L) Due to non-specific total JNK signal, band intensity was generated from solely normalized phosphorylated JNK to normalized alpha-tubulin. All lanes and data points represent biological replicates. Data is shown as mean ± standard error. Statistical significance was determined using one-way ANOVA with Dunnett correction (B, D, F, H, L) or Kruskal-Wallis tests with Dunn’s correction (J).

Western blot analysis of underlying, non-epidermal tissues in homeostatic vs. contralateral limbs indicated significantly decreased p38 phosphorylation in contralateral underlying tissues at 7 dpa compared to homeostatic non-epidermal tissues (p adj = 0.0276, Fig. 4I-J). The contralateral decrease in JNK phosphorylation was less pronounced in non-epidermal tissues, compared to the previously described epidermal comparisons, with solely the 7 dpa contralateral vs. homeostatic comparison retaining significance (p adj = 0.0345, Fig. 4K-L).

Interestingly, the contralateral decrease in ERK phosphorylation that was present in epidermal comparisons was not present in non-epidermal tissue comparisons (Fig. 4G-H). These results indicate that the long-range decrease in ERK phosphorylation induced by amputation is specific to the epidermis.

### Modification of MAPK signaling induces opposite changes in planarian epidermal permeability in homeostatic and amputation contexts

After observing that long-range epidermal permeability increases were associated with downregulated MAPK signaling in axolotls, we tested whether similar relationships were conserved between evolutionarily distant regenerative organisms. Specifically, we tested the effect of MAPK inhibition and activation on epidermal permeability in planarians, a commonly used model of regeneration (reviewed in Ivankovic et al., 2019; Sánchez Alvarado, 2012).

In DMSO-treated groups—where MAPK signaling was not manipulated—we found that amputated planarians unexpectedly exhibited a decrease in epidermal permeability, compared to homeostatic planarians (p adj = 0.0105, 0.0074, Fig. 5E, I). Amputation-induced decreases in planarian epidermal permeability were also recapitulated in DMSO-absent planarian water, with the effect persisting throughout the full 14-day regeneration period (all p adj < 0.0001, Fig. S2C).

**Fig 5.**
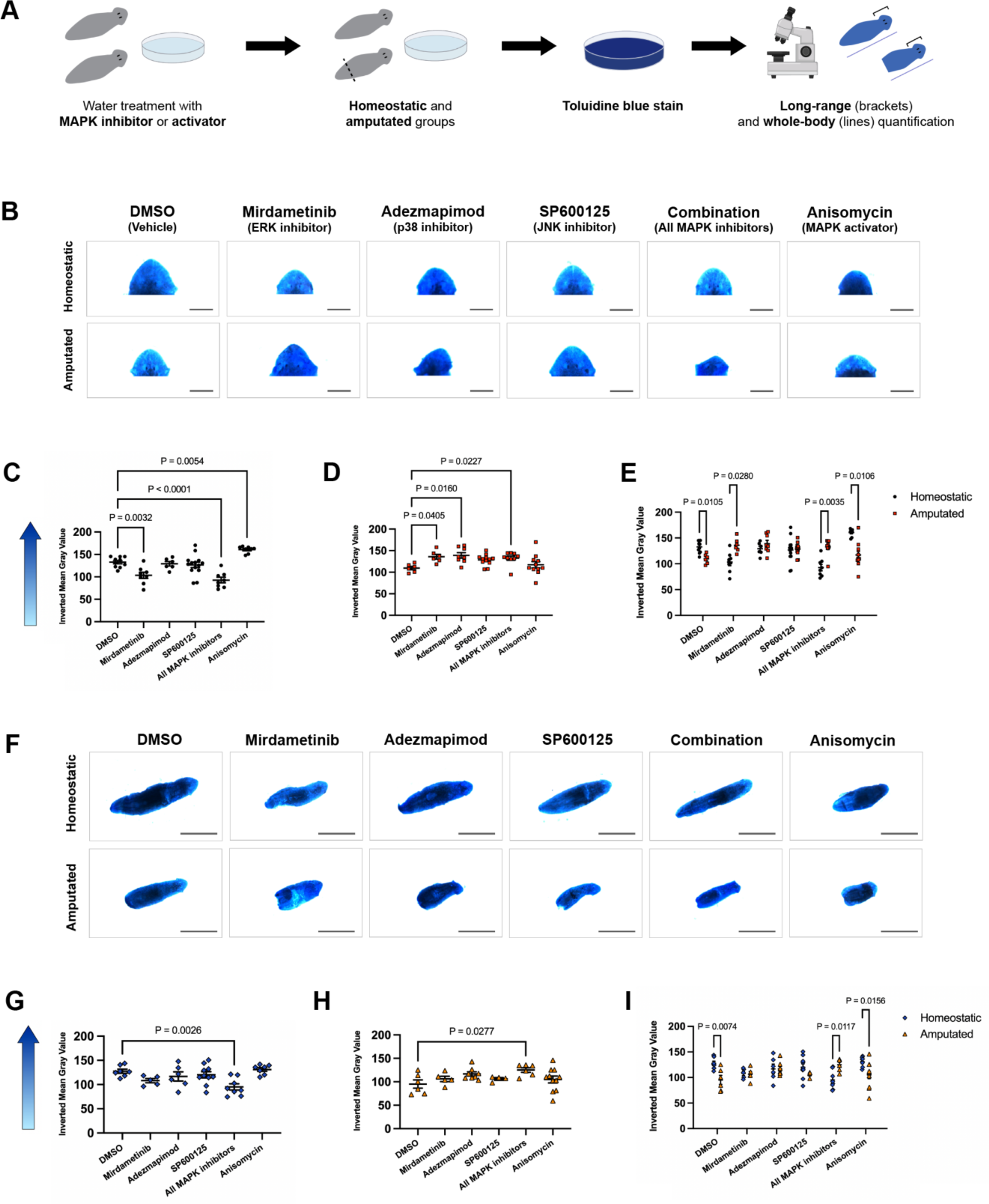
Modification of MAPK signaling induces opposite changes in planarian epidermal permeability in homeostatic and amputation contexts. (A) Experimental setup. (B) Representative images of stained planarian heads. (C-E) Quantification of long-range epidermal permeability in head tissue in (C) homeostatic and (D) amputation contexts, with (E) MAPK-dependent comparisons. (F) Representative images of stained whole planarians. (G-I) Quantification of whole-body epidermal permeability in (G) homeostatic and (H) amputation contexts, with (I) MAPK-dependent comparisons. Scale bars are (B) 0.5 mm or (F) 2 mm. Higher data points indicate darker blue staining and increased epidermal permeability. All data points represent biological replicates. Data is shown as mean ± standard error. Statistical significance was determined using Kruskal-Wallis ANOVA with Dunn’s correction (D), ordinary one-way ANOVA with Dunnett correction (C, G, H), Mann-Whitney U tests with Bonferroni-Dunn correction (E), or two-way ANOVA with Sidak correction (I).

We quantified epidermal permeability in the head of homeostatic planarians to match the tissue region used for analyzing “long-range” epidermal permeability in regenerating planarians.

Among non-amputated, homeostatic animals, we found that the epidermis of ERK-inhibited (p adj = 0.0032) and full-MAPK-inhibited (p adj < 0.0001) planarians was significantly less permeable than that of control animals (DMSO vehicle), where full-MAPK-inhibited planarians were simultaneously administered a combination of ERK, p38, and JNK inhibitors (Fig. 5C).

Conversely, the epidermis of uninjured planarians with activated MAPK signaling was significantly more permeable than that of DMSO control animals (p adj = 0.0054, Fig. 5C). These results indicate that in homeostatic uninjured planarians, inhibition of MAPK signaling is sufficient and necessary to decrease epidermal permeability (Fig. 5B-C).

However, when we looked at epidermal permeability in regenerating, as opposed to uninjured planarians, we found the opposite effect. The “long-range” epidermis (defined as the epidermis of the head, distant from the tail amputation plane) of ERK-inhibited (p adj = 0.0405), p38- inhibited (p adj = 0.016), and full-MAPK-inhibited (p adj = 0.0227) regenerating planarians was significantly more permeable than that of control regenerating planarians (DMSO vehicle, Fig. 5D), mirroring the link between decreased MAPK signaling and increased long-range epidermal permeability shown in regenerating axolotls. We found no significant difference in epidermal permeability between MAPK-activated and DMSO control regenerating planarians (Fig. 5B, D), possibly due to the intrinsic activation of MAPK signaling at baseline during planarian regeneration (Tasaki et al. 2011a; Tasaki et al. 2011b). These results indicate that the injury state of planarians—namely whether or not the animal is actively regenerating—modulates the effect of MAPK signaling on epidermal permeability.

Full-body analysis of epidermal permeability in planarians indicated similar, but less pronounced, trends as long-range epidermal analyses. The epidermis throughout the body of uninjured, full-MAPK-inhibited animals (p adj = 0.0026, Fig. 5G) was significantly less permeable than control animals (DMSO vehicle). Conversely, in regenerating planarians, the full-body epidermis of full-MAPK-inhibited animals was significantly more permeable than that of DMSO control planarians (p adj = 0.0277, Fig. 5H). Both long-range and full-body analyses highlighted that planarian amputation triggers contrasting changes in epidermal permeability, depending on levels of MAPK signaling (Fig. 5E, I).

## DISCUSSION

The findings of this study demonstrate a novel framework by which the combination of amputation and decreased MAPK signaling triggers an increase in epidermal permeability in areas distant from the wound site. We report epidermal permeability changes induced by amputation in two evolutionarily distant regenerative species, planarians and axolotls (Fig. 6).

**Fig 6.**
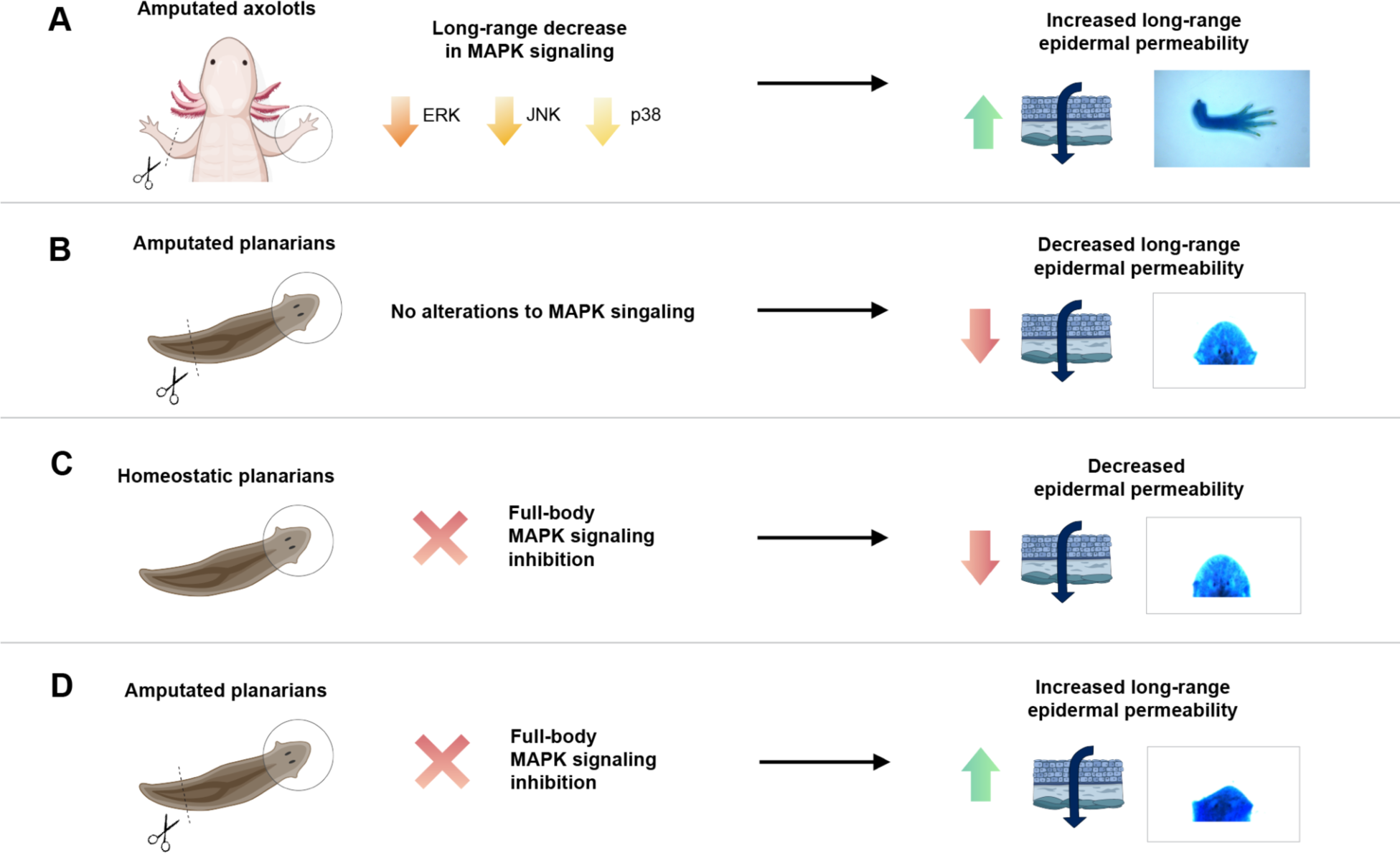
Summary of findings. An overview of the major findings of this study: (A) in axolotls, amputation triggers a long-range increase in epidermal permeability, accompanied by the downregulation of MAPK signaling in contralateral limbs; (B) in planarians, amputation triggers a long-range decrease in epidermal permeability; (C) in homeostatic planarians, full-body inhibition of MAPK signaling decreases epidermal permeability; and (D) in regenerating planarians, full-body inhibition of MAPK signaling increases long-range epidermal permeability.

In planarian regeneration, MAPK signaling mediates both localized and whole-body responses to amputation. ERK and JNK signaling regulate neoblast differentiation during blastema formation (Tasaki et al. 2011a; Tasaki et al. 2011b). Recently, ERK signaling was reported to mediate rapid transduction of whole-body, pro-regenerative signaling through planarian muscle tissue, highlighting the role of MAPK signaling in regulating systemic responses to amputation in planarians (Fan et al. 2023). MAPK signaling also promotes vertebrate regeneration (reviewed in Wen et al. 2022). ERK signaling, in particular, mediates wound healing after tail amputation in *Xenopus laevis* tadpoles (Sato et al. 2018), limb blastema formation in *X. laevis* froglets (Suzuki et al. 2007), antler regeneration in deer (Li et al. 2012), and fin and scale regeneration in zebrafish (De Simone et al. 2021; Owlarn et al. 2017).

Our results suggest that, beyond regeneration itself, regulation of MAPK signaling after amputation may also mediate epidermal barrier function. Indeed, MAPK signaling has been shown to play a role in epidermal function and structure in species other than axolotls and planarians. JNK1 deficiency has also been shown to impair barrier repair and wound healing in mice (Koehler et al. 2011), and p38 inhibition results in reduced expression of proteins involved in epidermal differentiation of human keratinocytes during homeostasis (Meng et al. 2018).

Furthermore, MAPK inhibition decreases filaggrin—a structural protein important for epidermal barrier maintenance—expression in human keratinocytes (Meng et al. 2018; reviewed in Sandilands et al. 2009). Given the shared utilization of MAPK signaling in maintaining epidermal barrier function, our findings in axolotls and planarians may indeed apply to non-regenerative models, including mammals, although further investigation of these questions is needed.

There are several possible explanations for the long-range downregulation of MAPK signaling during regeneration and its effects on epidermal permeability. In tissues distant from the zone of regeneration, pro-regenerative MAPK signaling may be decreased to prevent ectopic regeneration. While amputation induces body-wide cell cycle reentry in both axolotls and planarians (Johnson et al. 2018; Payzin-Dogru et al. 2024 preprint; Wenemoser and Reddien 2010), it is possible that blocking MAPK signaling in tissues distant from the amputation plane, prevents these primed cells from aberrantly converting to a full regeneration response. In this framework, a long-range increase in epidermal permeability may be a side effect, and possibly a vulnerability, of downregulating MAPK signaling in tissues distant from the zone of regeneration.

Conversely, there could be benefits to increasing epidermal permeability during regeneration in aquatic organisms like axolotls, most notably to promote respiration. One study reported that over half of the oxygen intake in the spotted salamander (*Ambystoma maculatum*) occurs through the epidermis (Whitford and Hutchison 1963), illustrating its importance for respiration. Metabolic state influences regeneration efficiency in the Iberian ribbed newt (*Pleurodeles waltl*) (Peng et al. 2021), and thus whole-body responses to increase oxygen consumption might be harnessed by salamanders to promote regeneration. There is also evidence that increased epidermal water permeability can be utilized in frog tadpoles as an anti-predator defense (Mori et al. 2009). These links suggest that in highly regenerative animals, changes in epidermal permeability could function to promote responses that maximize survival odds during vulnerable periods after amputation injury. Future assessment of the potential evolutionary functions of MAPK-signaling-regulated epidermal permeability changes is warranted to understand whether this phenomenon directly translates to mammals.

In mammals, dysregulation of MAPK signaling is known to recapitulate structural defects in inflammatory skin disorders like psoriasis and atopic dermatitis (Kanemaru et al. 2017).

Additionally, epidermal barrier dysfunction is relevant in amputation-related pathologies, including diabetes (Man et al. 2022). Amputees often experience dermatological side effects in the stump like contact dermatitis and infection (Koc et al. 2008; Levy 1980; reviewed in Lyon et al. 2000)—but systemic exacerbation or development of conditions like psoriasis beyond the wound site after amputation have also been reported, a progression possibly induced by whole-body stress responses to amputation (Heim et al. 2003). As such, investigating the relationship between MAPK signaling and epidermal barrier function, specifically in the context of amputation or multi-tissue injury, is clinically relevant.

Our study expands on a growing body of literature on the body-wide effects of amputation in regenerative organisms. We highlight previously uncharacterized functional changes in the epidermis during regeneration, by which amputation triggers long-range changes in epidermal permeability in both axolotls and planarians. Further study is needed to fully unravel the body-wide, evolutionarily-conserved consequences of amputation in the epidermis, which may guide approaches to therapeutics for amputation-related pathologies in humans.

## MATERIALS AND METHODS

### Animals

#### Axolotls

All animal experimentation was approved by and conducted in accordance with Harvard University’s Institutional Animal Care and Use Committee (Protocol #19-02-346). Age and size-matched leucistic axolotls (*Ambystoma mexicanum*) were used for all axolotl experiments.

Axolotls were housed in 40% Holtfreter’s Solution buffered to a salinity of 3000 microSiemens and pH of 7.6 and fed a diet of *Artemia*, rotifers, and/or Rangen soft-moist pellets depending on size. Axolotls were anesthetized in 0.1% tricaine prior to all experiments and allowed to recover overnight in 0.5% sulfamerazine after all surgical procedures.

#### Planarians

Planarians (*Dugesia dorotocephala*) were kept in water made with 0.5 g of instant ocean per 500 mL of deionized water. Planarians were fed 1 mL of *Artemia* per 30 animals once a week and water changed 5 and 24 hours after feeding. Planarians were fasted >7 days before amputation and were not fed during experimentation.

### MAPK inhibition and activation

#### Planarians

Planarians were kept in 10 μM mirdametinib (ERK inhibitor), 25 μM adezmapimod (p38 inhibitor), 25 μM SP600125 (JNK inhibitor), a combination of the three inhibitors (1 μM mirdametinib, 3 μM adezmapimod, 3 μM SP600125), 25 μM anisomycin (MAPK activator), or vehicle control (dimethyl sulfoxide, DMSO) water for 24 hours, based on established dosages (Bohr et al.; Tasaki et al. 2011a; Tasaki et al. 2011b; Xiong et al. 2006; Yuan et al. 2021).

Inhibitors were dissolved in 0.25% DMSO in planarian water by volume. After this treatment period, half of the planarians underwent a tail amputation at a distance of 1/4 of the total length from the tip of the tail. All animals were kept in their respective inhibitor, activator, or control treatment for a further 48 hours, then stained and imaged. Inhibitor details are listed in Table S1.

### Toluidine blue staining, quantification, and analysis

#### Axolotl limbs

Initial forelimb amputations were performed mid-zeugopod and the bone was trimmed back from the amputation plane. Amputations of collected limbs were performed mid-stylopod. Samples were dehydrated immediately after amputation for 2 minutes each in 25%, 50%, 75%, then 100% methanol at 4°C. Samples were then immediately rehydrated for 2 minutes each in 75%, 50%, 25% methanol and distilled water at 4°C then rinsed in 1X PBS for 2 minutes. Prepped samples were fully immersed in 0.1% toluidine blue solution (Sigma-Aldrich #89640-5G) for 2 minutes then de-stained in 1X PBS for 2 days at room temperature. Stained limbs were then imaged using a Leica M165 FC equipped with Leica DFC310 FX camera, and the mean gray value of each limb was quantified using a custom macro in ImageJ. For both axolotl and planarian quantifications, mean gray values were inverted using the formula Y’ = Y_max_ + Y_min_ - Y for ease of data interpretation on graphs. This transformation did not alter statistical findings.

For Fig. 1C, all groups were normal and had equal variances. One significant outlier was identified but included in the data as its inclusion did not change statistical findings; as such, statistical significance was determined using one-way ANOVA with Tukey’s HSD. For Fig. 1D, all groups had equal variances. Two groups within the data set (1 dpa and 70 dpa) were not normal; as such, statistical significance was determined using a non-parametric Kruskal-Wallis test with Dunn’s test of multiple comparisons. For Fig. 1E, all groups had equal variances. Two groups (1 dpa and 70 dpa) were not normal; as such, statistical significance was determined using non-parametric Mann-Whitney U tests with Holm correction.

For Fig. 1F, the segmenting methodology for regrowth vs. stump tissue is illustrated in Fig. 1G. All groups met normality, equal variance, and outlier assumptions; as such, statistical significance was determined using paired two-way ANOVA with Sidak correction.

#### Planarians

The same staining procedure was applied to planarians with two changes—planarians were dyed in 0.01% instead of 0.1% toluidine blue solution and de-stained in 1X PBS for one day at room temperature instead of two. Stained planarians were imaged using a Leica M165 FC equipped with Leica DF310 FX camera and selections for mean gray value calculations for each planarian were made manually.

For long-range measurements, planarian heads were quantified to measure epidermal permeability in tissues distant from the amputation plane. Full-body planarian selections were also quantified to measure body-wide changes in epidermal permeability.

For Fig. 5C, 5G, and 5H, groups met normality and equal variance assumptions; as such, statistical significance was determined using ordinary one-way ANOVA with Dunnett correction. For Fig. 5D, due to non-normality, statistical significance was determined using non-parametric Kruskal-Wallis ANOVA with Dunn’s correction. For Fig. 5E, due to non-normality, statistical significance was determined using non-parametric Mann-Whitney U tests with Bonferroni-Dunn correction. Groups in Fig. 5I met normality, equal variance, and outlier assumptions; as such, statistical significance was determined using two-way ANOVA with Sidak correction.

### Cryosectioning & limb preparation

After amputation, limbs were soaked in 4% paraformaldehyde-PBS for one hour at room temperature. They were then kept overnight in a 30% sucrose solution at 4°C. Following the overnight soak, limbs were embedded in optimal cutting temperature (OCT) compound and stored at -80°C until sectioning. Limbs were sectioned at a thickness of 16 µm on a Leica CM 1950 cryotome.

### Hematoxylin & eosin (H&E) staining

#### Staining

Slides were stained using the following protocol from Harvard University Department of Stem Cell and Regenerative Biology’s Histology Core:

Slides were soaked in water for 2 minutes, then Harris Hematoxylin (Sigma #HHS-128) for 3 minutes, water for 2 minutes, quickly dipped twice in acid alcohol (5 mL HCl, 368 mL 95% ethanol, 127 mL water), then soaked in water for 1 minute. Slides were then stained in Scott’s Bluing (2 g potassium bicarbonate, 20 g magnesium sulfate in distilled water), soaked in water for 1 minute, stained in eosin (Fisher #23-314631) for 1 minute, then rinsed in 95% ethanol for 2 x 1 min, 100% ethanol for 3 x 1 min, and Histoclear for 3 x 1 min. Slides were then covered with a coverslip and imaged on a Zeiss Axio Scan.Z1 automated slide scanner.

#### Quantification

For each biological replicate, two 800 μm-long epidermal segments were selected and averaged from both anterior and posterior sides of the radial, carpal, and digit regions of the most representative full limb sections for quantification. Epidermal thickness was calculated via Fiji ImageJ and nuclear density was calculated via CellProfiler. For Fig. 2C and 2D, all groups met normality and equal variance assumptions; as such, statistical significance was determined using ordinary one-way ANOVA with Tukey correction. For Fig. 2E, all groups met normality, equal variance, and outlier assumptions; as such, statistical significance was determined using a paired two-way ANOVA with Sidak correction. For Fig. 2F, due to non-normality, statistical significance was determined using non-parametric Kruskal-Wallis ANOVA with Dunn’s correction.

### Immunofluorescence

#### Staining

For antibody staining, frozen sections were rehydrated with PBS at room temperature. Sections were blocked and permeabilized for 30 minutes at room temperature with 0.1% Triton X-100 in either 10% goat serum-PBS or 10% donkey serum-PBS, depending on the host of the secondary antibody. Sections were incubated with primary antibody overnight at 4°C. The following day, slides were washed with PBS for five minutes three times, after which sections were incubated with the appropriate secondary antibody for 2.5 hours at room temperature.

Sections were washed with PBS, incubated with DAPI, and washed with PBS once more before mounting. Slides were imaged on a Zeiss Axio Scan.Z1 automated slide scanner. Parameters for primary and secondary antibodies are listed in Table S1.

#### Claudin 1 and collagen IV quantification

Epidermal segments were selected from radial, carpal, and digit regions of full limb sections for quantification of mean fluorescence intensity. For claudin 1, these segments were defined as 800 μm-long with full epidermal thickness. Leydig cells were masked out of each selection by nuclear size and DAPI intensity thresholds. For collagen IV, these segments were defined as 300 μm-long and 30 μm-thick, with the basement membrane positioned centrally in each segment. Mean fluorescence intensity was calculated via CellProfiler by dividing the total intensity of each image selection by the area of tissue in each selection. For Fig. 3C, groups met normality, equal variance, and outlier assumptions; as such, statistical significance was determined using ordinary one-way ANOVA with Tukey correction. For Fig. 3D and 3H, groups met normality, equal variance, and outlier assumptions; as such, statistical significance was determined using a paired two-way ANOVA with Sidak correction. For 3G, both groups were normal; as such, statistical significance was determined using a two-tailed paired t-test.

### Western Blots

#### Protocol

Protein isolation and Western blotting were performed as previously described (Sousounis et al. 2020). One µg of protein per well was used for ERK blots and 2 ug of protein were used for JNK and p38 blots. PVDF 0.2 μm membranes (Bio Rad #1620177) were used instead of nitrocellulose membranes. After imaging non-loading-control targets, membranes were subsequently stripped with Restore PLUS stripping buffer (Thermo Scientific #PI46430) for 10 minutes, followed by loading control incubation and imaging, using the same protocol. Primary and secondary antibody parameters are listed in Table S1.

#### Quantification

Films were scanned at 600 dpi, with brightness and contrast adjusted equally across biological replicates within each row. Area under curve (AUC) values were quantified in Fiji ImageJ. AUC values were normalized within each row. Subsequently, AUC values for each protein target of interest were normalized to their respective AUC values for alpha tubulin loading control. For ERK and p38 blots, band intensity was then generated from the ratio of normalized phosphorylated to total protein AUC values. Due to non-specific total JNK signal, band intensity was generated from solely phosphorylated JNK to alpha-tubulin. For Fig. 4B, 4D, 4F, 4H, and 4L, all groups passed normality and equal variance assumptions; as such, statistical significance was determined using ordinary one-way ANOVA with Dunnett correction. For Fig. 4J, due to non-normality, statistical significance was determined using non-parametric Kruskal-Wallis ANOVA with Dunn’s correction.

## DATA AVAILABILITY STATEMENT

Data and code will be made available upon request.

## CONFLICT OF INTEREST

J.L.W. is a co-founder of Matice Biosciences. Other authors declare no conflicts of interest.

## ACKNOWLEDGEMENTS

We would like to thank the Histology Core of Harvard University’s Department of Stem Cell and Regenerative Biology for their contributions to the H&E stains. We would like to express our gratitude to Isaac Adatto, Damian Bernard, Brianna Blackmore, Nicholas Cardelia, Hayden Graham, Lauryn Wilson, Omenma Abengowe, Erin Anderson, Rui Qun Miao, and Vicky Yan for their assistance with animal care. We are grateful to members of the Whited Lab for their valuable advice and discussions during this study. We would also like to thank Amanda Flores at the Harvard TH Chan School of Public Health for her technical assistance with Adobe Illustrator.

## AUTHOR CONTRIBUTIONS

Conceptualization: K.E.D., T.D., D.P., J.L.W.; Methodology: K.E.D., R.T.K., E.M.K., E.C., J.B., T.D.; Software: A.A., B.T., K.E.D., R.T.K.; Validation: K.E.D., R.T.K., E.M.K., E.C.; Formal analysis: K.E.D., R.T.K., E.M.K., E.C., A.G.L., H.S., S.Y.C.W.; Investigation: K.E.D., R.T.K., E.M.K., E.C., N.L., C.J.P., K.T., A.S.; Resources: K.E.D., H.S., J.L.W.; Data curation: K.E.D., R.T.K., E.M.K., E.C., J.P., A.G.L., S.Y.C.W., H.S.; Writing - original draft: K.E.D., R.T.K., J.P., J.L.W.; Writing - review & editing: K.E.D., R.T.K., A.A., B.T., J.P., S.Y.C.W., D.P., J.L.W.; Visualization: K.E.D., R.T.K.; Project Administration: K.E.D., R.T.K.; Supervision: K.E.D., R.T.K., D.P., J.L.W.; Project administration: K.E.D., R.T.K., D.P., J.L.W.; Funding acquisition: R.T.K., J.L.W.

## FUNDING

This work was supported by the NSF-CAREER IOS-2145925 (J.L.W.) and NICHD R01HD115272 (J.L.W.), the Harvard Herchel Smith Undergraduate Science Research Program (R.T.K.), and the Harvard Program for Research in Science and Engineering (R.T.K.).

---

**Fig S1.**
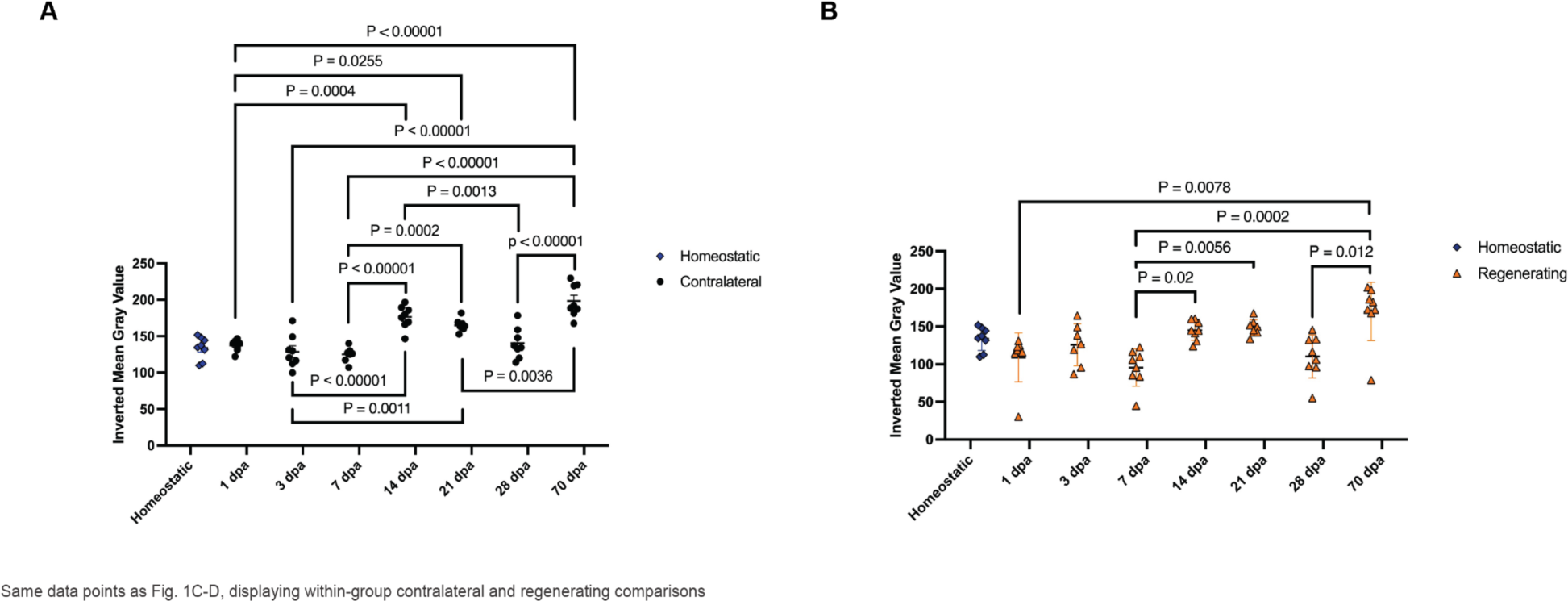
Epidermal permeability changes over time in contralateral and regenerating axolotl limbs. (A) Quantification of inverted mean gray values of homeostatic and contralateral limbs. Data points shown are the same as Fig. 1C, displaying inter-time point comparisons between contralateral groups. (B) Quantification of inverted mean gray values of homeostatic and regenerating limbs. Data points shown are the same as Fig. 1D, displaying inter-time point comparisons between regenerating groups. In both graphs, data is shown as mean ± standard error. Statistical significance was determined using one-way ANOVA with Tukey’s HSD (A) and Kruskal-Wallis ANOVA with Dunn’s test of multiple comparisons (B).

**Fig S2.**
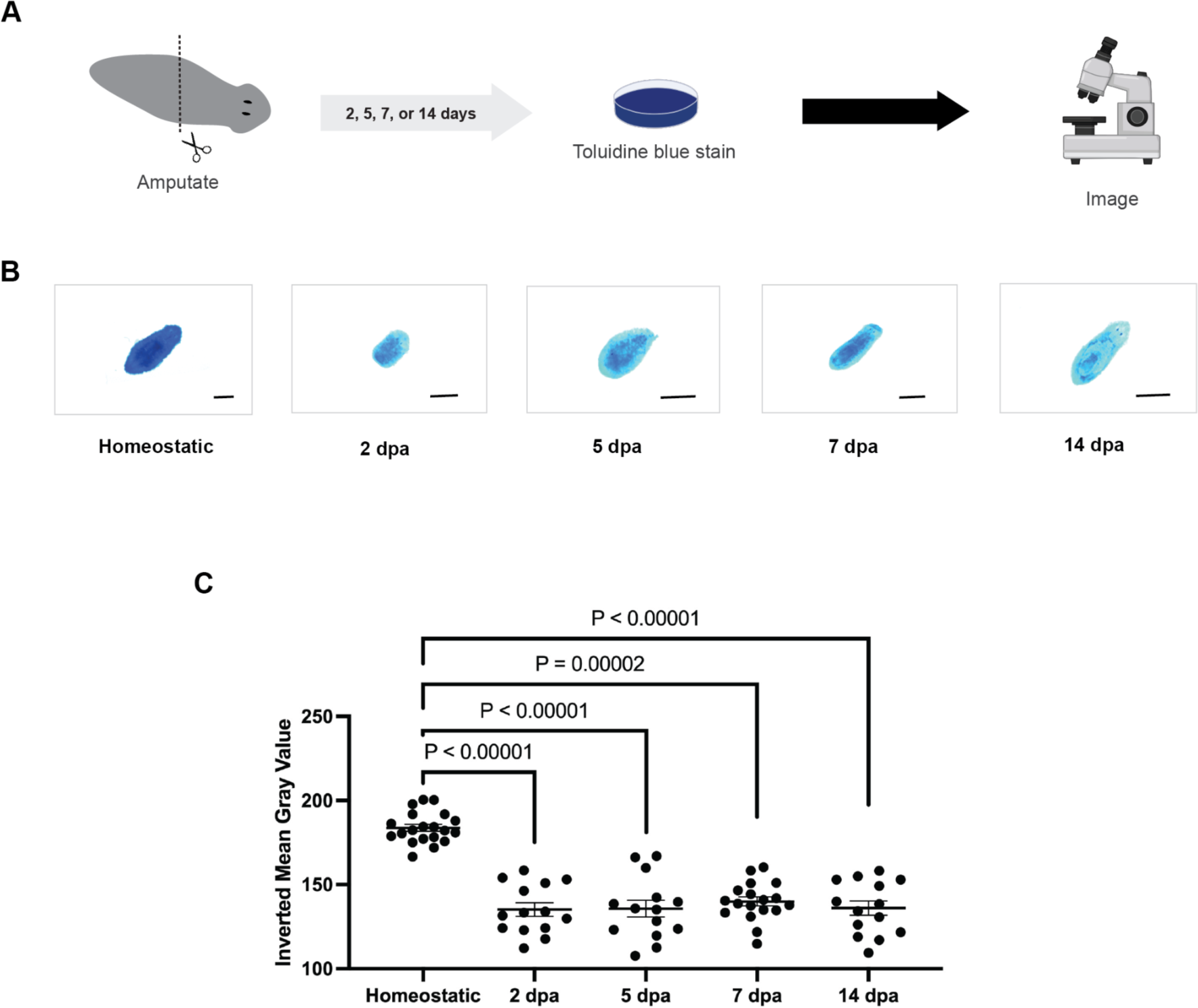
Amputation triggers decreased epidermal permeability in planarians that persist throughout regeneration. (A) Experimental set-up. (B) Representative images of stained planarians. Bars are 1 mm. (C) Quantification of inverted mean gray values. Higher inverted mean gray values indicate darker planarians. Data is shown as mean ± standard error. Statistical significance was determined using Kruskal-Wallis ANOVA with Dunn’s test of multiple comparisons.

**Table S1.** Key resources.

| REAGENT or RESOURCE | SOURCE | IDENTIFIER |
| --- | --- | --- |
| Primary antibodies |  |  |
| Rabbit anti-phospho-ERK1/2, Thr202/Tyr204 (WB 1:500) | Cell Signaling | Cat#: 4370 |
| Rabbit anti-ERK1/2 (WB 1:500) | Cell Signaling | Cat#: 9102 |
| Rabbit anti-phospho-SAPK/JNK, Thr183/Tyr185 (WB 1:250) | Cell Signaling | Cat#: 9251 |
| Rabbit anti-SAPK/JNK (WB 1:250) | Cell Signaling | Cat#: 9252 |
| Rabbit anti-phospho-p38 MAPK, Thr180/Tyr182 (WB 1:250) | Cell Signaling | Cat#: 9211 |
| Rabbit anti-p38 MAPK (WB 1:250) | Cell Signaling | Cat#: 9212 |
| Rabbit anti-claudin 1 (IF 1:100) | Thermo Fisher | Cat#: 71-7800 |
| Rabbit anti-collagen IV (IF 1:100) | Abcam | Cat#: ab6586 |
| Secondary antibodies |  |  |
| Goat anti-rabbit HRP (WB 1:5000) | Bio-Rad | Cat#: 1721019 |
| Goat anti-mouse HRP (WB 1:5000) | Jackson ImmunoResearch | Cat#: 115-035-003 |
| Donkey anti-rabbit 488 (IF 1:100) | Jackson ImmunoResearch | Cat#: 711-545-152 |
| Goat anti-rabbit Cy3 (IF 1:500) | Jackson ImmunoResearch | Cat#: 111-165-003 |
| Chemicals |  |  |
| Mirdametinib/PD0325901 (ERK inhibitor) | MedChemExpress | Cat#: HY-10254 |
| Adezmapimod/SB203580 (p38 inhibitor) | Selleck Chemicals | Cat#: S1076 |
| SP600125 (JNK inhibitor) | MedChemExpress | Cat#: HY-12041 |
| Anisomycin (MAPK activator) | MedChem Express | Cat#: HY-18982 |

## Notes

### Summary of Updates

We clarify which species of planarians was used, Dugesia dorotocephala.

